# Disentangling competitive and cooperative components of the interactions between microbial species

**DOI:** 10.1101/2024.02.06.579244

**Authors:** Aamir Faisal Ansari, Gayathri Sambamoorthy, Thrisha C Alexander, Yugandhar B.S. Reddy, Janhavi Raut, Narendra M. Dixit

**Affiliations:** Department of Chemical Engineering, Indian Institute of Science, Bengaluru, India; Unilever R&D India Pvt Ltd, Bengaluru, India; Department of Bioengineering, Indian Institute of Science, Bengaluru, India

**Author notes:** Correspondence: Narendra M. Dixit.

## Abstract

Interactions between microbial species have been characterized by the net influences, positive or negative, that each species in a pair exerts on the other. This conventional view of interactions being either positive or negative proves restrictive in predicting the behaviour of microbial communities and, more importantly, influencing them towards desired community structures. Here, we propose a more fundamental characterization of the interactions. The net interactions typically comprise positive and negative underlying components. Yet, the conventional view prevails because the components have been difficult to disentangle. We have developed a methodology to disentangle them when metabolic interactions predominate. We conceived a theoretical resource partitioning between species that helps estimate the positive components. The negative components then follow from knowledge of the net interactions. The interactions between two species are then characterized by the ‘quartet’ of these components. We applied the methodology *in silico* to 28 species pairs from the human oral microbiome, yielding 56 net interactions and their 112 quartet components. We found that on average the net interactions comprised positive and negative components of comparable strengths. Interestingly, weak net interactions often arose from the cancellation of strong underlying components. Furthermore, we found species pairs with similar net interactions but vastly different underlying components. Extant community ecology theories, based on net interactions, cannot distinguish between such pairs. The quartet explained several confounding experimental observations and offered new insights into microbial community ecology. We envision its implications in the construction of more refined ecological theories and the engineering of synthetic microbial communities.

## INTRODUCTION

Multi-species microbial communities are important to human health (1), the environment (2), and biotechnology (3). Establishing and maintaining stable microbial communities relies on the knowledge of the interactions between the constituent species (4). The interactions have proven challenging to unravel and understand (5-11). Conventionally, the interactions between species pairs have been categorized based on the nature–positive or negative–of the effect each species has on the survival and/or growth of the other: When the effects are both negative (-,-), *i*.*e*., when the species inhibit each other, the species are said to exhibit competition. When the effects are both positive (+,+), the species are said to exhibit cooperation, or mutualism. Other combinations are denoted exploitation or parasitism (+,-), amensalism (-,0), commensalism (+,0), and neutralism (0,0), where ‘0’ implies no effect. A question under active investigation today is whether the interactions between species pairs are predominantly competitive or cooperative (5, 7, 10, 12-21). On one side, evidence exists that cooperative interactions improve the efficiency of the microbial community, by distributing the metabolic burden between species or enabling new collective functionalities (13-15). Cooperation has been seen in several settings; for instance, it occurs between species in the gut microbiome and between species grown under extreme, resource starved conditions (16, 17). On the other side, ecological theory argues that cooperative interactions may introduce dependencies that may render communities unstable (10). Competitive interactions, particularly exploitative ones like the predator-prey interactions, are predicted to be stabilizing as they introduce negative feedbacks. In agreement, experiments on culturable bacteria from several niches have found a predominance of competitive interactions between species (5, 7, 12, 18-21). A comprehensive understanding of the factors determining the predominance of competition versus cooperation is still lacking.

Here, we argue that the prevalent ‘either-or’ view of competition and cooperation may be restrictive in describing species interactions. A more nuanced view that accounts for the simultaneous presence of competitive and cooperative interactions is warranted. The latter view is prompted by the observation, made in several studies, of species pairs exhibiting either competition or cooperation depending on environmental conditions (8, 17, 22-24). For instance, increasing resource availability or the species diversity in a community altered the interactions between species pairs in the community from cooperation to competition (8, 17, 23). Thus, species pairs appear capable of engaging in both competitive and cooperative interactions. The origins of competition and cooperation are typically distinct. When interactions are predominantly metabolic in nature, typical of many ecological niches and synthetic communities (23, 25-27), competition arises from shared resources, whereas cooperation is attributed to cross-feeding metabolites. Thus, competitive and cooperative interactions are not exclusive; they can occur together. Accordingly, we hypothesized that a species pair would engage ‘simultaneously’ in competitive and cooperative interactions and that the observed interactions would be the ‘net’ effect of these opposing components. Depending on which component dominates, the net interaction would manifest as competition or cooperation. The dominant component may be determined by the specific environmental conditions, including resource availability. Knowledge of the competitive and cooperative components would offer a deeper, more fundamental understanding of species interactions than the net interactions alone. For instance, two species pairs with similar net interactions may respond quite differently to environmental changes if the net interactions were made up of vastly different underlying components in the two cases. Accounting for the components may thus lead to more refined ecological theories and help better engineer microbial communities. While the net interactions are readily measured for culturable species (e.g., (5, 7, 17, 22, 24)), no methods exist for disentangling their competitive and cooperative components.

Here, we devised a method to estimate the competitive and cooperative components of the interactions between species pairs. Assuming a predominance of metabolic interactions, we constructed a theoretical partitioning of resources that enables the estimation of the cooperative component needed to maintain one of the species in the community when the other is sustained entirely by the resources. The competitive components are then estimated from knowledge of the net interactions. Species pairs can then be defined by a quartet of interactions: the cooperative and competitive components of the interaction of each species with the other.

## RESULTS

### Method to estimate the cooperative and competitive components of species interactions

We consider a two-species community, comprising species ‘1’ and ‘2’, in an environment supplied with nutrients (or resources) at a constant rate (Fig. 1A). We denote the steady-state growth rates of the species in the environment as 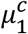 and 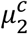, respectively, where the superscript ‘c’ refers to the community. In the same environment, we denote the steady-state growth rates of the species when present alone, or in monoculture, as 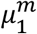 and 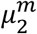, respectively. If the species experienced no interactions, the monoculture and community growth rates of the respective species would be identical. A difference between the two rates would indicate the presence of net interactions between the species (Fig. 1B and 1C). Following convention, we defined the net influence of species ‘2’ on species ‘1’ as:

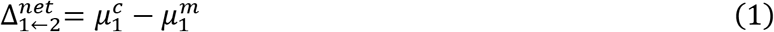

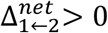 would imply an overall positive effect of species 2 on species 1, whereas 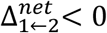 would imply a net negative effect. Similarly, we defined the net influence of species ‘1’ on species ‘2’ as

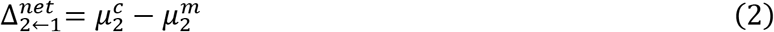

Note that the pair of net interactions 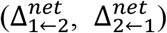 is conventionally used to classify the nature of the interactions between species pairs, which we summarized above. Our goal is to estimate the positive and negative components of these net interactions.

**Figure 1.**
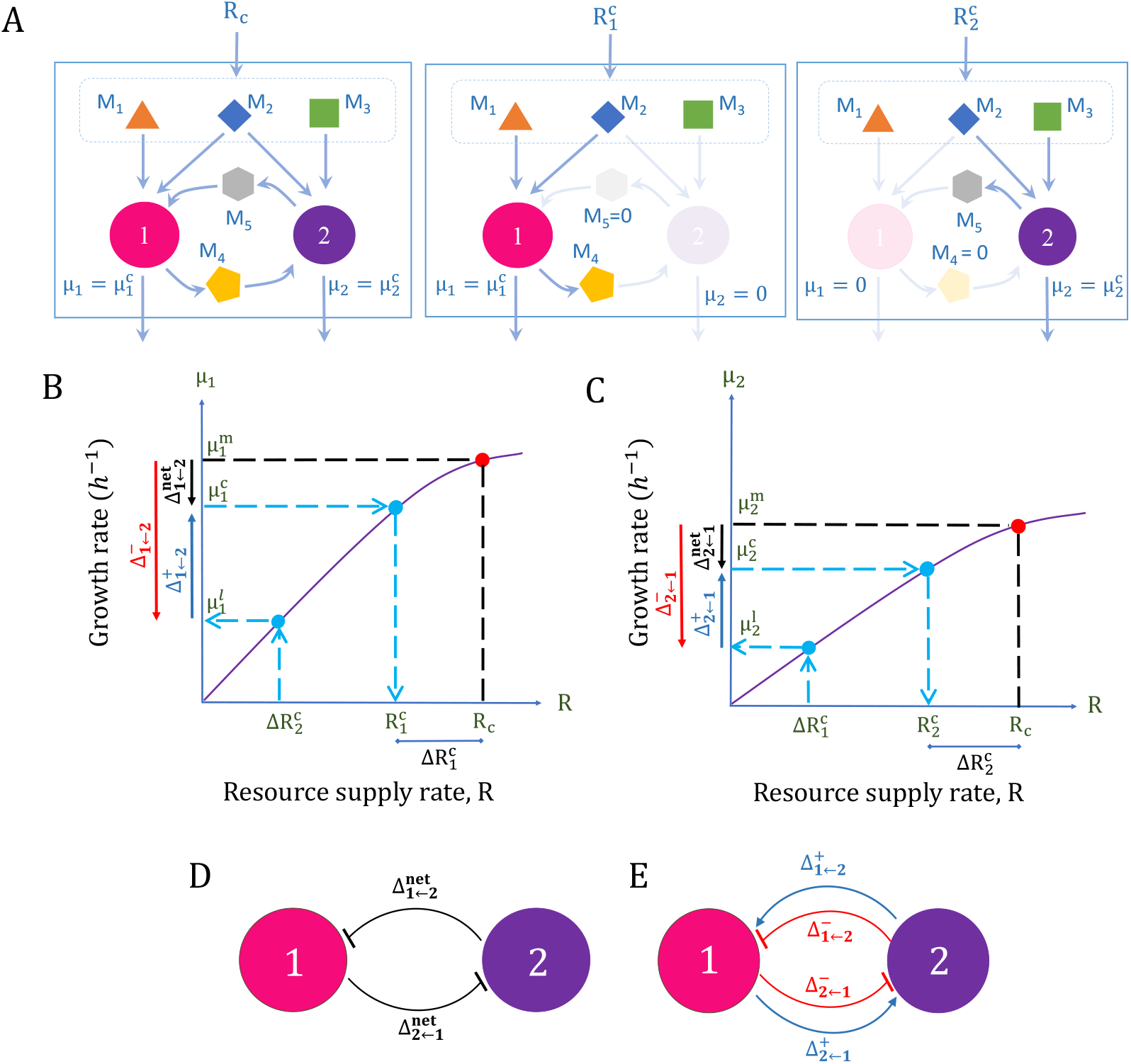
Illustration of our method to estimate the positive and negative components of the net interactions. (A) A hypothetical two-species community supplied with nutrients, M1, M2, and M3, at constant rates, with *R*_*c*_ the supply rate of the shared limiting resource, M2 (left). Metabolite M4, produced by species 1 (pink) and consumed by species 2 (purple), and metabolite M5, with the opposite pattern, are cross-feeding metabolites. The species growth rates in the community are 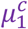 and 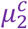, respectively. The resource supply rates that support the species individually at their community growth rates are 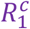 (middle) and 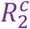 (right). (B & C) The growth-resource curves of species 1 (B) and species 2 (C) in monoculture. 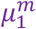 and 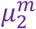 are the monoculture growth rates of the species at the resource supply rate *Rc*. The leftover resources, 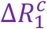 and 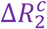, yield the associated growth rates, 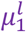 and 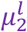. The interaction components estimated by our method (see text) are illustrated next to the y-axis. (D) A depiction of the net interactions employed conventionally. (E) A depiction of the quartet of components estimated here.

The method we devised is based on the following conceptual arguments. When species 1 and 2 are both present in a community, resources are expected to be partitioned between them, resulting in competitive interactions. The specific partitioning, however, is not typically known (22, 23, 28), precluding easy quantification of the competitive components. We conceived of a theoretical partitioning that would yield a bound on the competitive component and thereby offer an estimate of the cooperative component. Accordingly, we let species 2 receive all the resources necessary for sustaining its growth at the rate observed in the community. The ‘left-over’ resources would then be available for use by species 1. Using knowledge of how the growth of species 1 depends on resource availability, we estimate the growth rate of species 1 that the left-over resources could sustain. If the latter growth rate is smaller than the observed growth rate of species 1, then it would imply that species 1 is sustained additionally by cross-feeding metabolites from species 2 in the community. Indeed, the difference between the observed growth rate and the growth rate from the left-over resources would be an estimate of the positive component of the influence of species 2 on species 1. In the same way, with another theoretical partitioning where species 1 gets all the resources needed to sustain its growth in the community, one can estimate the positive component of the net influence species 1 has on species 2. The difference between the net influence, estimated above, and the positive component would yield the negative component. Below, we formalized these conceptual arguments into a procedure for estimating the components.

We denoted the supply rate of the limiting resource in the environment as *R*. The growth rates of the species are expected to increase monotonically with *R*. We defined the curve depicting the growth rate, *μ*, of a species in monoculture as a function of *R* as the ‘growth-resource curve’ of the species. This curve is readily determined using experiments, or using genome-scale metabolic models where available, and is analogous to the classical description of the growth rate dependent on the substrate concentration due to Monod (29). We employed the growth-resource curves of the two species (Fig. 1B and 1C) to estimate the positive components.

We let *R*_*c*_ be the resource supply rate to the community. Note that this is the supply rate at which the growth-rates of the species in the community were 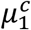 and 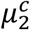 and in monocultures were 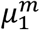 and 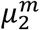, respectively, as mentioned above. Using the growth-resource curve of species 2, we estimated the resource supply rate, 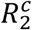, at which the growth rate of the species 2 in monoculture would equal its growth rate in the community, 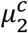 (Fig. 1C). If 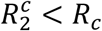, it implied that not all of the resources supplied to the community were needed for the maintenance of species 2. The remaining, or left-over, limiting resource supply rate, 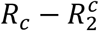, denoted 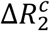, would therefore be available to species 1 (Fig. 1C). We now used the growth-resource curve of species 1 and identified its growth rate in monoculture afforded by the left-over resource supply rate 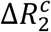, which we denoted as 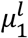 (Fig. 1B). 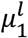 would thus be the growth rate of species 1 that the environment would sustain ‘after’ letting species 2 exist at its community growth rate in the absence of additional interactions between the two species. Clearly, the difference between 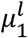 and the observed growth rate, 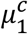, must be due to the positive effect of species 2 on species 1. We thus estimated the positive component of the interaction as

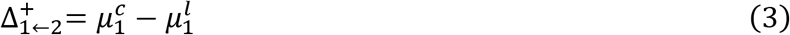

The difference between the net interaction, estimated above (Eq. 1), and the positive component yielded the negative component:

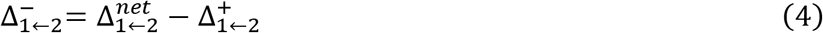

In the same way, we estimated the components for the effect of species 1 on species 2: We employed the growth-resource curve of species 1 to obtain the resource supply rate, 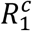, at which its monoculture growth rate would equal 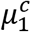 (Fig. 1B). The left-over resource 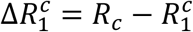 would be available to species 2 after letting species 1 grow at its community growth rate, 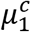. From the growth-resource curve of species 2 (Fig. 1C), the growth rate, 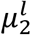, of species 2 at the left-over resource supply rate 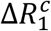 was estimated. The positive component of the net influence of species 1 on species 2 was then

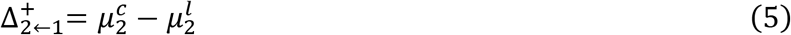

and the negative omponent,

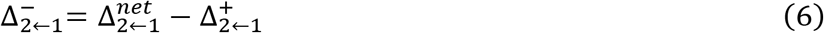

Together, the components 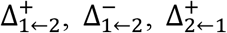, and 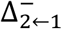 formed the quartet of interactions between the two species in the community environment (Fig. 1D, E). Below, we demonstrate the procedure by applying it to a specific species pair.

### The quartet of interactions between *Actinomyces viscosus* and *Bifidobacterium lactis*

We considered the species *Actinomyces viscosus* (Av) and *Bifidobacterium lactis* (Bf), which are part of the human oral microbiome (30). Using genome-scale metabolic reconstructions of the two species (Methods), we predicted their individual growth-resource curves. We considered resources resembling a Western diet and quantified the resource supply rate, *R*, as the fold-increase in the supply rate of the limiting resources over a basal rate (Methods). Over the range of *R* we examined, the growth-resource curves were linear (Fig. 2A and B). At any *R*, the growth rate of Av, *μ*_*Av*_ was higher than that of Bf, *μ*_*Bf*_. For instance, at *R* = 1, the rates were: *μ*_*Av*_ ∼0.27 *h*^*−*1^ and *μ*_*Bf*_ ∼0.04 *h*^*−*1^.

**Figure 2.**
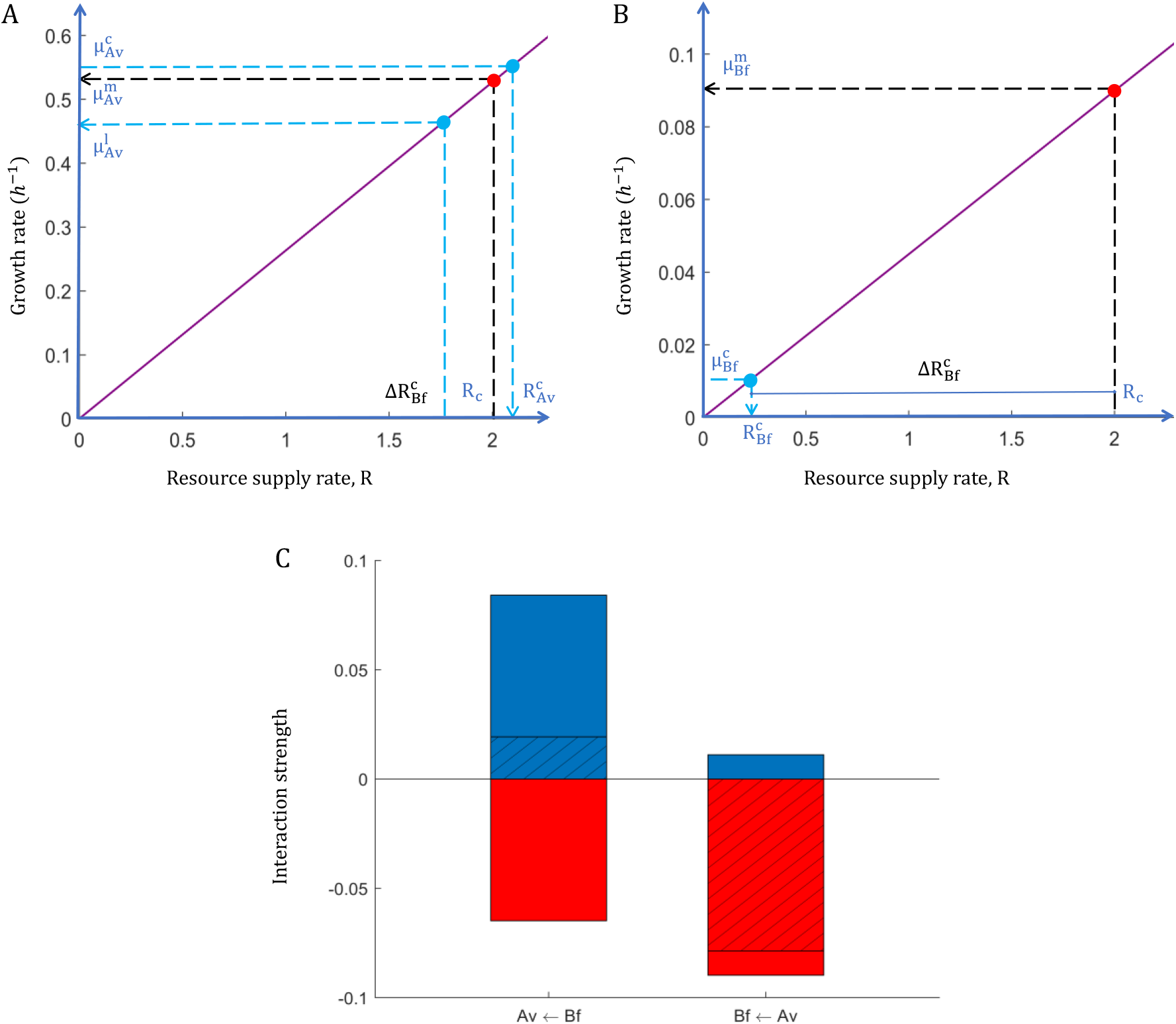
Components of interactions between the species Av and Bf. The growth-resource curves of Av (A) and Bf (B) and the community, monoculture and left-over growth rates of the two species indicated (see text). (C) The positive (blue) and negative (red) interaction components, forming the interaction quartet, and the net (black hatched) interactions between the species.

We now considered a community environment with a resource supply rate *R*_*c*_ = 2 and estimated the monoculture and community growth rates of the species (Methods). The growth rate of Av in the coculture was 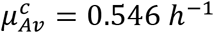, while that of Bf was 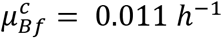. The corresponding monoculture growth rates of the species were 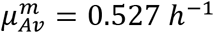 and 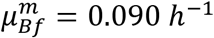. From these estimates, we obtained the net interactions as the difference of the growth rates in the coculture and monoculture, so that 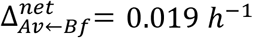 and 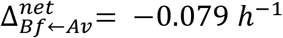 (Fig. 2C). The pair thus exhibited exploitative interactions, where Av gained from Bf but Bf suffered due to Av.

To estimate the components of these net interactions, we identified the resource supply rate required for each species to sustain its community growth rate in the absence of additional interactions. Thus, for Av, the resource supply rate 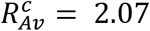 (Fig. 2A) ensured 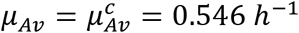, whereas for Bf the resource supply rate needed for sustaining *μ*_*Bf*_ at 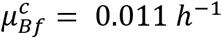 was 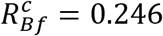 (Fig. 2B). Because *R*_*c*_ = 2 is smaller than 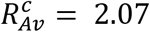, sustaining Av at its community growth rate would not leave any resource for Bf; *i*.*e*., 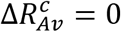 (Fig. 2A). On the other hand, the resource left over after sustaining Bf at its community growth rate would be 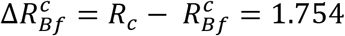. We estimated the growth rate of Av that can be afforded by the latter left-over resource to be 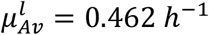 (Fig. 2B). The positive components of the interactions were thus 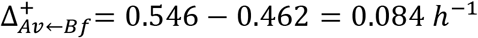 and 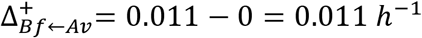. The negative components followed as the differences between the net interactions and the positive components: 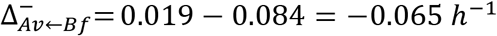 and 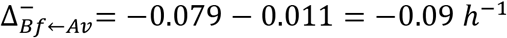. Our method thus yield the quartet for the Av-Bf pair: 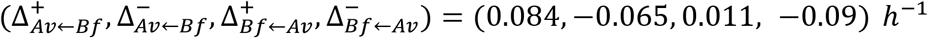 (Fig. 2C). Following this procedure, we next estimated the quartets of a large number of species pairs in the human oral microbiome.

### The quartets of interactions between species in a representative oral microbiome

We considered 8 species that have been employed previously to construct a synthetic human oral microbiome (9, 31): *Streptococcus mutans* (Smu), *Streptococcus mitis* (Smi), *Streptococcus parasanguinis* (Sp), *Streptococcus salivarius* (Sl), *Streptococcus sanguinis* (Ss), and *Lactobacillus casei* (Lc), along with Bf and Av. We first estimated the net pair-wise interactions between the species. We employed genome-scale metabolic models of each of the species and estimated the monoculture and community growth rates of the species in all possible 2-species communities (Methods; Tables S1-S7). We employed two different methods, SteadyCom (32) and MICOM (33), to estimate the growth rates in the two species communities, and to ensure that our findings were not specific to the community modelling method. We present the results obtained using SteadyCom here. The results with MICOM, which were qualitatively similar to those with SteadyCom, are in the Supplementary Materials (see Text S1, Fig. S1, Tables S5-S7). Among the 28 species pairs, resulting in 56 net interactions, we found that 29 interactions were net positive and 27 interactions were net negative (Fig. 3A). The strengths of the positive and negative interactions were comparable: the median positive interaction strength was 0.14 *h*^*−*1^ (IQR: [0.06 0.20] *h*^*−*1^), whereas the median negative interaction strength was *−*0.15 *h*^*−*1^ (IQR: [*−*0.11 *−*0.66] *h*^*−*1^) (Fig. 3A). Interestingly, of the 28 species pairs, 27 pairs showed exploitation or parasitic interactions, while only a single pair exhibited mutualism (the Sp-Lc pair) (Fig. 3C). This predominance of exploitation/competition and the limited occurrence of mutualism is consistent with ecological theory and measurements on culturable bacterial species (7, 10, 12, 18-21).

**Figure 3.**
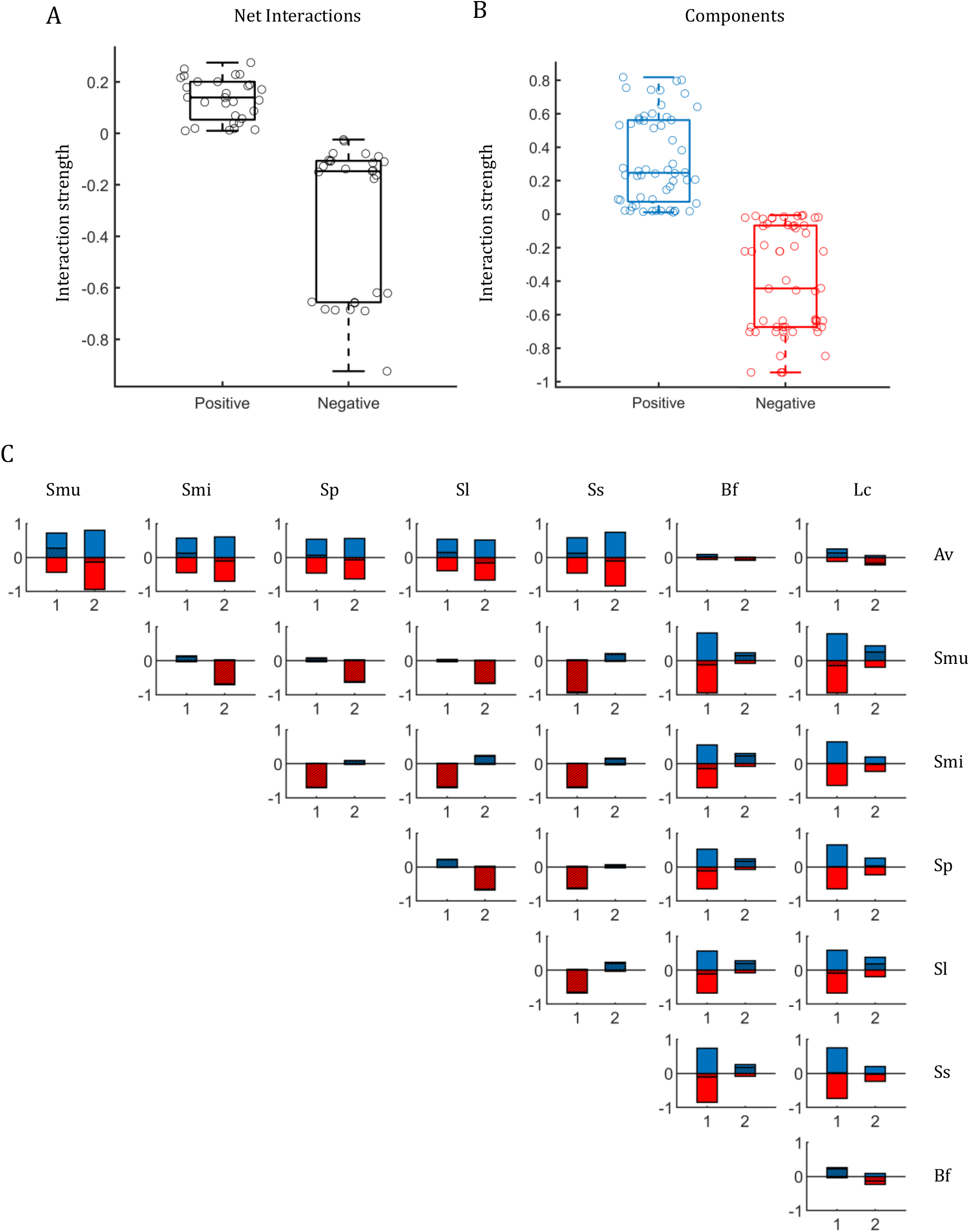
Estimation of the quartet of interactions for species pairs from a representative oral microbiome. The distribution of (A) the net interactions and (B) their components between all pairs of 8 species in a representative human oral microbiome (see text). In (A), the net positive and net negative interactions are shown separately for clarity. (C) The individual interactions (hatched black) and their components (blue – positive; red – negative). In each panel, 1 is the species mentioned next to the row (row species) and 2 that above the column (column species) to which the panel belongs. The x-axis labels (1 or 2) indicate the species on which the influence of the other (2 or 1) is estimated. In (A) and (B), boxes show median and interquartile ranges and whiskers show extremes.

We next estimated the components of these net interactions using our method above (Fig. 3B and 3C). We exploited the linear dependence of the growth rates on the resource supply rate (see Fig. 2A) to estimate the components without explicitly computing the entire growth-resource curves (Methods). Of the 112 components (constituting the 56 net interactions), we found 106 to be non-zero, indicating that the net interactions were, in a vast majority of the cases, the resultant of significant underlying components (Fig. 3B and 3C). Overall, the positive and negative components estimated were equally powerful: The median positive component was 0.25 *h*^*−*1^ (IQR: [0.08 0.56] *h*^*−*1^) and the median negative component was *−*0.45 *h*^*−*1^ (IQR: [*−*0.07 *−*0.68] *h*^*−*1^) (Fig. 3B). The latter ranges were comparable to the ranges of the net interactions above.

Importantly, positive components existed in several cases where the net interactions were negative. For instance, Av had a net negative influence on Ss, with 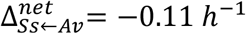. However, on estimating the quartet, we found that this small net negative interaction was made up of large positive and negative components: 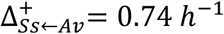 and 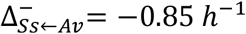. Similarly, negative components existed when the net interactions were positive. In the same species pair, we found that Ss had a positive effect on Av, with 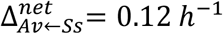, and that the components of the latter interaction were 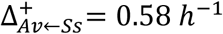, and 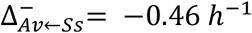. The net interactions, positive or negative, between species pairs were thus the resultants of the underlying positive and negative components.

Further, interestingly, we found cases where the net interactions between species pairs were similar but were made up of vastly different underlying components. For instance, the Av-Smu pair had similar net interactions to the Bf-Lc pair: Specifically, 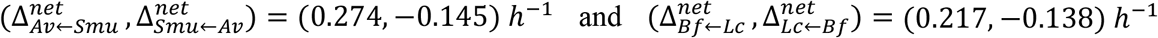. The components, however, were much larger in magnitude for the Av-Smu pair than the Bf-Lc pair. The quartets were: 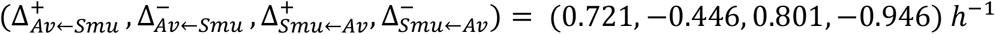 and 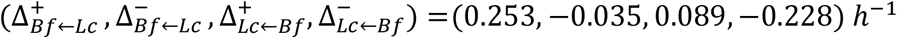. It is possible, therefore, that the two pairs will behave differently when subjected to the same environmental changes, a difference that extant theories reliant on the net interactions alone would not capture.

The set of the components, or the quartet, is thus a more fundamental representation of species interactions than the net interactions. We examined next the implications of the knowledge of the quartet.

### Insights into the behaviours of microbial communities

Varying environmental conditions has been argued to alter the interactions between species (17, 22-24). Broadly, in the conventional view based on net interactions, increasing resource availability is argued to minimize the need for cooperation, thereby increasing competition between species pairs (6, 12, 17). As resource availability increases, however, the production of cross-feeding metabolites may increase as well, as they are products of the metabolism of the resources, thereby potentially increasing the extent of cooperation. How cooperation diminishes with increasing resources is thus not fully clear. Here, to address this question, we examined how the quartet of interactions between two species varied with resource availability. We considered the pair of Av and Bf, examined above. Recall that at *R*_*c*_ = 2, the pair exhibited exploitation. We now varied *R*_*c*_ over a wider range, from 0 to 6, and estimated the variation of the quartet.

We found, interestingly, that all the components increased monotonically with *R*_*c*_. Indeed, consistent with the expectation above, as *R*_*c*_ increased from 1 to 2, the positive component 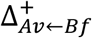 increased more than its negative counterpart 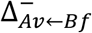, so that the net interaction of Bf on Av became more positive (Fig. 4). Beyond 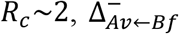 increased more steeply with *R*_*c*_ than 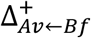. At, 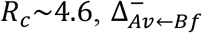 exceeded 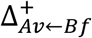, so that the net influence of Bf on Av switched from being positive to negative. Increasing *R*_*c*_ further increased the extent of the negative interaction. Throughout, the influence of Av on Bf stayed negative (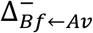 was larger than 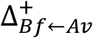) and increased its magnitude with *R*_*c*_. Thus, the Av-Bf pair switched from exploitation to competition upon increasing *R*_*c*_ beyond ∼4.6. The latter transition is consistent with the expectation of increasing competition with resource availability. The decomposition of the net interactions into the quartet helps understand this transition consistently with the expectation of increasing cross-feeding metabolites with resources.

**Figure 4.**
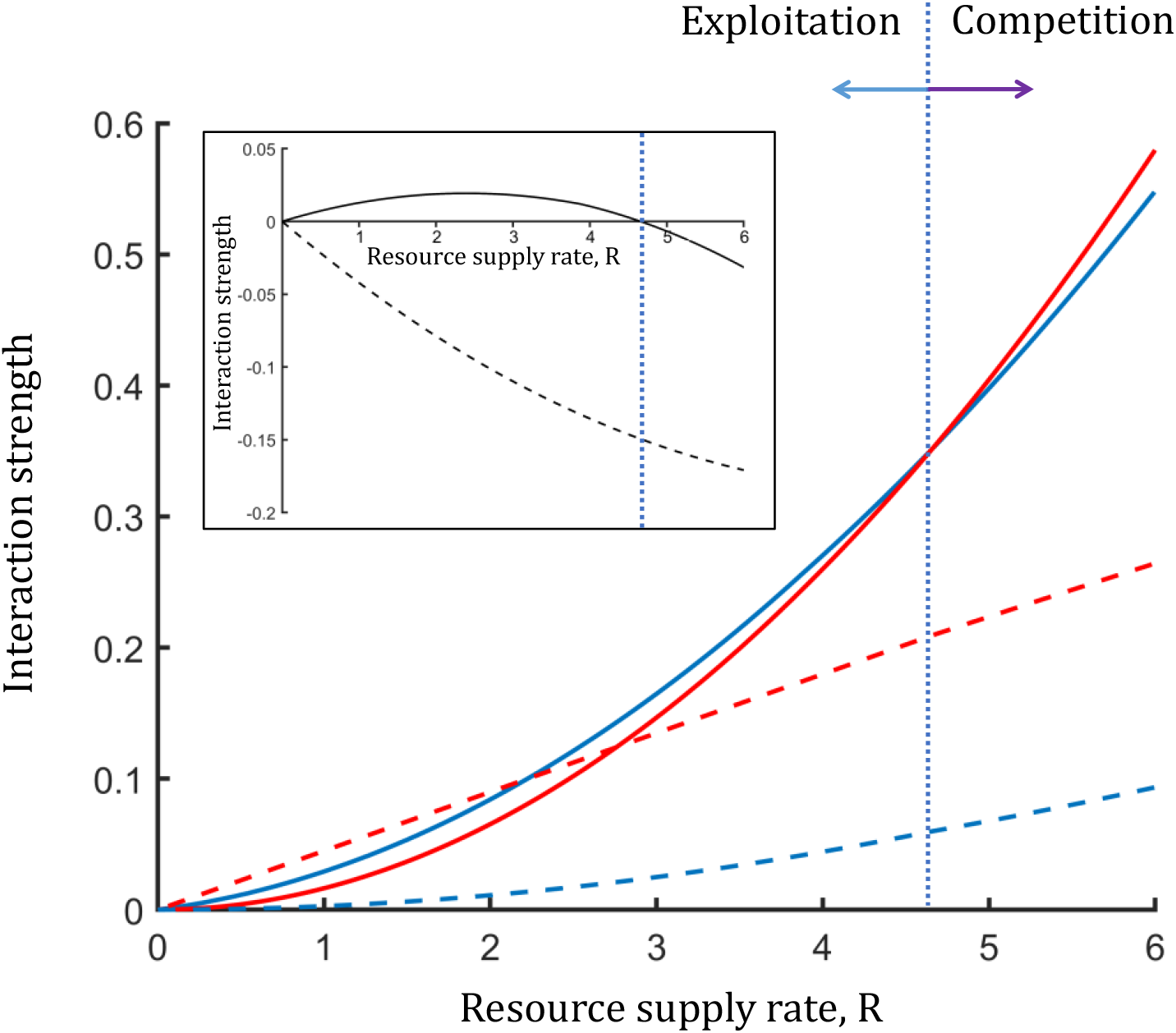
Variation of the quartet with resource availability for the Av-Bf pair. The positive (blue) and negative (red) components of the influence of Bf on Av (solid lines) and Av on Bf (dashed lines) as a function of the resource supply rate, showing the transition from exploitation (left of blue dotted line) to competition (right of blue dotted line). *Inset*: The corresponding net interactions.

Further implications follow from the predictions above for understanding the stability of communities. Current ecological theories predict that weak interactions are stabilizing (10, 17). While this may be true in general, the transition from cooperation to competition with increasing resource availability, a major change, must occur in a region in which the net interactions vanish (they change signs) and are thus necessarily weak. Our predictions suggest a reconciliation. We recognize that while the net interactions indeed vanish across such a transition, the components need not; rather, they may remain large across the transition. In our calculations above, it is the differential variation of the components with resource availability that underlies the transitions (Fig. 4). The weak net interactions are thus a result of the cancellation of the effects of potentially strong opposing components and may thus not necessarily be stabilizing. Future theories and experiments may test the robustness of these observations.

## DISCUSSION

The prevalent paradigm of categorizing species interactions using the overall, or net, influence species have on each other has resulted in major advances in our understanding of microbial community ecology (10). The paradigm, however, has limitations. Growing evidence argues for a more nuanced view of species interactions, given that species may engage in both competitive and cooperative interactions (8, 17, 22-24). Here, we proposed a more fundamental representation of species interactions that relies on a decomposition of the overall interactions into their competitive and cooperative parts, yielding a quartet of interactions governing species pairs. We developed an algorithmic methodology for estimating the quartet. Important insights emerged when we applied the methodology *in silico* to species pairs from a representative human oral microbiome.

We found that a vast majority of the interactions were made up of both cooperative and competitive components. Thus, positive components existed when species exhibited net negative interactions and vice versa. Furthermore, we found that weak net interactions could be made up of strong underlying components. Extant ecological theories, which are based on the net interactions alone, have argued that weak interactions are stabilizing (10, 34). Whether the predictions hold when the weak net interactions arise from a cancellation of strong underlying components is not clear. It is possible that perturbations to the strong underlying components may manifest as significant changes, including changes of sign, in the net interactions, potentially compromising community stability. Future theories that explicitly incorporate the quartet may help establish more robust criteria for community stability.

Interestingly, we found species pairs that had comparable net interactions but that were made up of vastly different underlying components. This finding again highlights limitations of extant ecological theories that are based on net interactions (10, 34, 35). The latter theories would predict that the two species pairs would respond similarly to environmental changes. One would expect, however, that the pairs would respond differently. For instance, increasing resource availability is expected to obviate the need for cooperative interactions (12, 17, 23). We speculate that the pair with stronger cooperative components may be affected more by increasing resource availability than the other pair. Stronger cooperative components may reflect a greater inadequacy of the supplied resources, which increasing resource availability may alleviate. Future experiments may test these predictions.

Knowledge of the quartet offers a more fundamental standpoint than the net interactions to understand the behaviour of microbial communities. To illustrate this, we examined the impact of increasing resource availability on a two species community. Although increasing resource availability may diminish the need for cooperation (in agreement with extant theories), in scenarios where cross-feeding metabolites drive cooperation, the opposite effect may ensue. Increasing resources may increase the production of cross-feeding metabolites and potentially enhance cooperation. Indeed, our calculations using genome-scale metabolic models of species did predict increased cooperation with increasing resources. At the same time, it predicted a switch to net competition when resources were increased beyond a threshold (Fig. 4), consistent with extant theories. The reconciliation between these conflicting observations came from knowledge of the quartet. With increasing resources, both positive and negative components increased, but the latter increased more than the former, resulting in the eventual switch in the net interaction type. We recognize that the switch may not always occur. If, for instance, the positive component continued to increase with resource availability more than the negative component, greater cooperation would occur upon increasing resource availability. Indeed, in recent experiments, species pairs that experienced limited overlap in the utilization of resources and therefore experienced restricted competition, saw an increase in cooperation with increasing resource availability (24). Costless metabolites, which incur minimal production costs but can be cross-fed (36), may also increase cooperation with increasing resource availability. Incorporating the quartet may thus facilitate the construction of more refined ecological theories and improve our ability to design and engineer synthetic microbial communities for applications in biotechnology and healthcare.

Our study has limitations. First, it is restricted to scenarios dominated by metabolic interactions. While such interactions are the more common ones (23, 26, 27, 37), scenarios where other types of interactions, driven for instance by toxins or signalling molecules (38), dominate may find behaviours distinct from that predicted by our study. Second, our method yields upper bounds on the magnitudes of the components. By allotting one of the species all the resources necessary for its sustenance in the community and letting the other species be sustained by resources and positive interactions, the latter interactions may be overestimated. Third, our study has relied on *in silico* analysis using genome-scale models, which may not capture the true interactions between species (23, 31). For instance, genome-scale models correctly predicted observed shifts between negative and positive interactions between species upon changing media conditions in only 65% of the cases (23). Our predictions are thus to be viewed as conceptual rather than quantitative. Nonetheless, we ascertained the robustness of our findings by performing our analysis using two different community modelling approaches, which yielded qualitatively consistent results. Besides, our methodology to identify the quartet can be implemented experimentally: The components can be estimated from the growth-resource curves of the species involved and their community growth rates, which are typically easily measured. Finally, our study is restricted to pair-wise interactions, whereas high-order interactions may exist in multispecies communities (8, 9, 31, 39). Future studies may consider developing the application of the quartet to the latter settings.

## METHODS

### Species metabolic models

We applied our method to species pairs from a representative oral microbial community (31). The community contains *Streptococcus mutans* (Smu), *Streptococcus mitis* (Smi), *Streptococcus parasanguinis* (Sp), *Streptococcus salivarius* (Sl), *Streptococcus sanguinis* (Ss), *Lactobacillus casei* (Lc), *Bifidobacterium lactis* (Bf), and *Actinomyces viscosus* (Av). We retrieved their metabolic models from the Virtual Metabolic Human database (40, 41) (VMH database), which were curated semi-automatically. These reconstructions are state-of-the-art, publicly available metabolic models for the microbes we considered. We set all ATP maintenance requirements, which are not known for these species, to zero. We used the createMultiSpeciesModel and SteadyCom functions of the COBRA Toolbox (42) to create and analyze metabolic models of cocultures, respectively. MICOM was implemented in Python.

The nutrient medium we selected followed the Western diet (40). We identified the set of common exchange reactions between the species involved and let it constitute the base medium for the species. To make the nutrient medium richer or poorer relative to the base medium, we multiplied the maximum uptake rates in the base medium by a fixed factor. For instance, when the factor was 2, the medium had maximum uptake rates twice those in the base medium.

Microbiomes usually exist under balanced growth conditions (32), which is characterized by equal specific growth rates of all species, also called the community growth rate, *μc*. SteadyCom guarantees the balanced growth of species in the community (32). The method employs the nutrient supply rates and computes the community growth rate, *μc*, and the species compositions. From the latter, we estimated the growth rates of the individual species in the community as follows. If *x*_*i*_ were the relative abundance of species *i*, then its ‘fractional’ growth rate, or its relative contribution to the community growth rate, would be *μ*_*c*_*x*_*i*_, which we used as a proxy for the species growth rate, 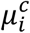. MICOM also considers balanced growth but estimates the growth rates, 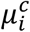, using an alternative heuristic approach that leads to realistic growth rate estimates in multispecies communities (33).

### Estimation of interaction components

The relationship between the growth rate, *μ*, and the limiting resource supply rate, *R*, is typically linear. (The saturation of the growth rate in Monod kinetics (29) implies that the resource is no longer limiting.) Capitalizing on this linearity, we estimated the quartet of interactions without explicitly constructing the growth-resource curves. We first estimated the monoculture and community growth rates under the given resource supply setting. To estimate left-over growth rates, we defined the ‘pure competition curve’ as the locus of points in the *μ*_1_ vs. *μ*_2_ plot where the interactions were purely competitive, *i*.*e*., no cooperative interactions existed (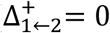 and 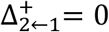). By definition, on this curve, 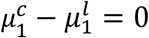 and 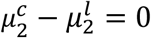. We denoted 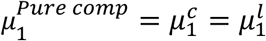 and 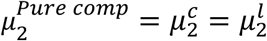. The set of points 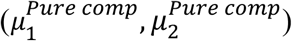 thus defined the pure competition curve.

We first obtained the pure competition curve for the case where *μ*_1_ = *a*_1_*R* and *μ*_2_ = *a*_2_*R*. We estimated 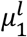 and 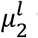 using our method: 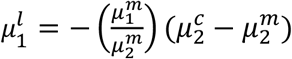 and 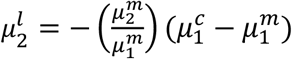. Using the above definition, the pure competition curve is thus given by: 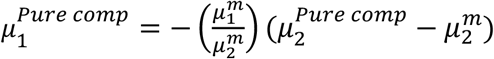 and 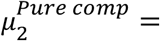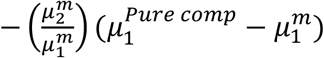. In other words, the pure competition curve for this case is the straight line passing through 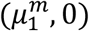 and 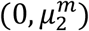 in the *μ*_1_ vs. *μ*_2_ plot. The intersection of the line 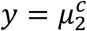 with the pure competition line yields 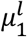 as its *x*-coordinate. Similarly, the intersection of the line 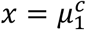 with the pure competition line yields 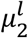 as its *y* coordinate. The interactions then follow as described in Fig. 2A. If the linearity of the dependence of the growth rate on resource supply rates is violated, then the full growth-resource curves would have to be estimated. The procedure outlined in the main text for the estimation of the components then follow.

## ACKNOWLEDGEMENTS

We thank Karthik Raman for his comments. GS acknowledges support from the IISc-IoE program.

