## Supplementary Materials for "Disentangling competitive and cooperative components of the interactions between microbial species"

**Contents:**

Text S1. Estimates of the quartets using MICOM

Figure S1. Estimation of the quartet of interactions for species pairs from a representative oral microbiome based on growth rates calculated using MICOM.

Table S1. Growth rates of species in monocultures.

Table S2. Growth rates of species in pairwise cocultures calculated using SteadyCom.

Table S3. Net interactions between species estimated based on growth rates calculated using SteadyCom.

Table S4. The quartets of the interactions between species estimated based on growth rates calculated using SteadyCom.

Table S5. Growth rates of species in pairwise cocultures calculated using MICOM.

Table S6. Net interactions between species estimated based on growth rates calculated using MICOM.

Table S7. The quartets of the interactions between species estimated based on growth rates calculated using MICOM.

**Text S1. Estimates of the quartets using MICOM**

Here, we summarize our findings of the estimation of the quartets using MICOM for community modelling. We recall that with SteadyCom, among the 28 species pairs we examined, resulting in 56 net interactions, 29 interactions were net positive and 27 interactions were net negative (main text, Fig. 3). Further, of the 28 species pairs, 27 pairs showed exploitation or parasitic interactions, while a single pair exhibited mutualism. Using MICOM, we found 24 net positive (median $0.32\ h^{-1}$; IQR: $[0.18, 0.44]\ h^{-1}$), and 32 net negative interactions (median $-0.19\ h^{-1}$; IQR: $[-0.12, -0.31]\ h^{-1}$), with 14 pairs exhibiting exploitative interactions, 9 competitive interactions, and 5 mutualistic interactions (Fig. S1). Thus, the overall predominance of exploitative and competitive interactions and the low prevalence of mutualistic interactions was consistent with the findings with SteadyCom. Further, of the 112 components (constituting the 56 net interactions), we had found 106 to be non-zero with SteadyCom. With MICOM, we found that all the components were non-zero. The median positive component was $0.49\ h^{-1}$ (IQR: $[0.24, 0.71]\ h^{-1}$) and the median negative component was $-0.47\ h^{-1}$ (IQR: $[-0.23, -0.67]\ h^{-1}$) (Fig. S1). The latter were comparable to the estimates from SteadyCom (main text, Fig. 3).

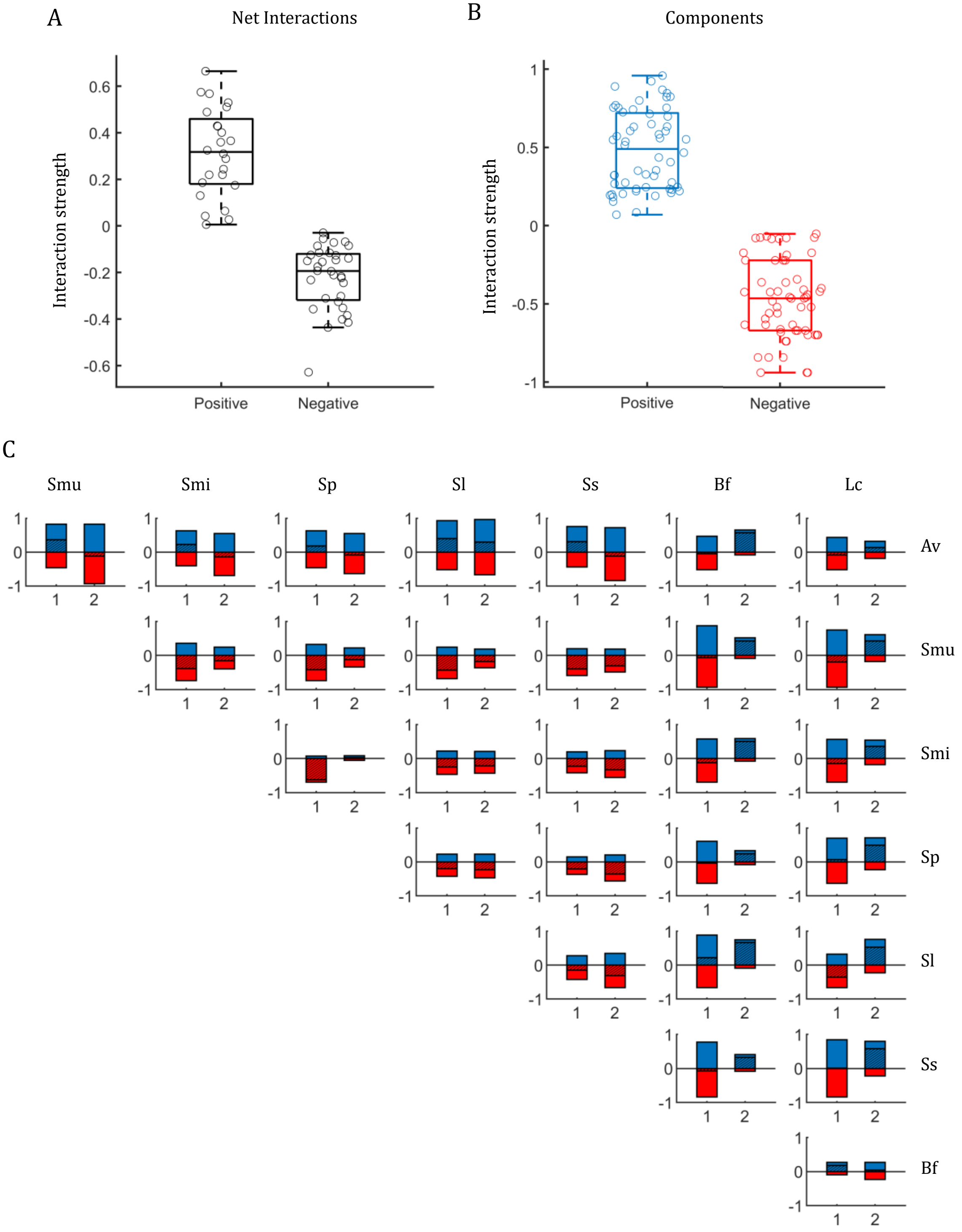


**Figure S1. Estimation of the quartet of interactions for species pairs from a representative oral microbiome based on growth rates calculated using MICOM.** The distribution of (A) the net interactions and (B) their components between all pairs of 8 species in a representative human oral microbiome (see text). In (A), the net positive and net negative interactions are shown separately for clarity. (C) The individual interactions (hatched black) and their components (blue – positive; red – negative). In each panel, 1 is the species mentioned next to the row (row species) and 2 that above the column (column species) to which the panel belongs. The x-axis labels (1 or 2) indicate the species on which the influence of the other (2 or 1) is estimated. In (A) and (B), boxes show median and interquartile ranges and whiskers show extremes.

**Table S1. Growth rates of species in monocultures.** Units are $h^{-1}$.

| Av | Smu | Smi | Sp | Sl | Ss | Bf | Lc |
|---|---|---|---|---|---|---|---|
| 0.522 | 0.937 | 0.698 | 0.634 | 0.670 | 0.841 | 0.089 | 0.226 |

**Table S2. Growth rates of species in pairwise cocultures calculated using SteadyCom.** Subscript 1 refers to the species in the row and 2 to the column. Units are $h^{-1}$.

| | Smu | | Smi | | Sp | | Sl | | Ss | | Bf | | Lc | |
|---|---|---|---|---|---|---|---|---|---|---|---|---|---|---|
| | $\mu_1^c$ | $\mu_2^c$ | $\mu_1^c$ | $\mu_2^c$ | $\mu_1^c$ | $\mu_2^c$ | $\mu_1^c$ | $\mu_2^c$ | $\mu_1^c$ | $\mu_2^c$ | $\mu_1^c$ | $\mu_2^c$ | $\mu_1^c$ | $\mu_2^c$ |
| **Av** | 0.801 | 0.801 | 0.650 | 0.601 | 0.595 | 0.562 | 0.666 | 0.514 | 0.648 | 0.739 | 0.546 | 0.011 | 0.655 | 0.052 |
| **Smu** | | | 1.062 | 0.022 | 1.002 | 0.021 | 0.959 | 0.020 | 0.021 | 1.038 | 0.818 | 0.245 | 0.796 | 0.478 |
| **Smi** | | | | | 0.015 | 0.727 | 0.018 | 0.900 | 0.020 | 0.988 | 0.558 | 0.319 | 0.715 | 0.204 |
| **Sp** | | | | | | | 0.856 | 0.018 | 0.018 | 0.890 | 0.529 | 0.260 | 0.652 | 0.267 |
| **Sl** | | | | | | | | | 0.021 | 1.049 | 0.562 | 0.290 | 0.587 | 0.412 |
| **Ss** | | | | | | | | | | | 0.743 | 0.269 | 0.867 | 0.198 |
| **Bf** | | | | | | | | | | | | | 0.307 | 0.090 |

**Table S3. Net interactions between species estimated based on growth rates calculated using SteadyCom.** Subscript 1 refers to the species in the row and 2 to the column. Units are $h^{-1}$.

| | Smu | | Smi | | Sp | | Sl | | Ss | | Bf | | Lc | |
|---|---|---|---|---|---|---|---|---|---|---|---|---|---|---|
| | $\Delta_{1\leftarrow 2}^{net}$ | $\Delta_{2\leftarrow 1}^{net}$ | $\Delta_{1\leftarrow 2}^{net}$ | $\Delta_{2\leftarrow 1}^{net}$ | $\Delta_{1\leftarrow 2}^{net}$ | $\Delta_{2\leftarrow 1}^{net}$ | $\Delta_{1\leftarrow 2}^{net}$ | $\Delta_{2\leftarrow 1}^{net}$ | $\Delta_{1\leftarrow 2}^{net}$ | $\Delta_{2\leftarrow 1}^{net}$ | $\Delta_{1\leftarrow 2}^{net}$ | $\Delta_{2\leftarrow 1}^{net}$ | $\Delta_{1\leftarrow 2}^{net}$ | $\Delta_{2\leftarrow 1}^{net}$ |
| **Av** | 0.274 | -0.145 | 0.123 | -0.105 | 0.068 | -0.078 | 0.139 | -0.163 | 0.121 | -0.110 | 0.019 | -0.079 | 0.128 | -0.176 |
| **Smu** | | | 0.116 | -0.684 | 0.056 | -0.620 | 0.013 | -0.657 | -0.925 | 0.188 | -0.129 | 0.155 | -0.150 | 0.250 |
| **Smi** | | | | | -0.691 | 0.086 | -0.687 | 0.223 | -0.685 | 0.139 | -0.148 | 0.229 | 0.009 | -0.024 |
| **Sp** | | | | | | | 0.216 | -0.659 | -0.622 | 0.040 | -0.111 | 0.170 | 0.012 | 0.039 |
| **Sl** | | | | | | | | | -0.656 | 0.200 | -0.115 | 0.200 | -0.090 | 0.184 |
| **Ss** | | | | | | | | | | | -0.107 | 0.179 | 0.018 | -0.030 |
| **Bf** | | | | | | | | | | | | | 0.217 | -0.138 |

**Table S4. The quartets of the interactions between species estimated based on growth rates calculated using SteadyCom.** Subscript 1 refers to the species in the row and 2 to the column. Units are $h^{-1}$.

| | Smu | | | | Smi | | | | Sp | | | | Sl | | | | Ss | | | | Bf | | | | Lc | | | |
|---|---|---|---|---|---|---|---|---|---|---|---|---|---|---|---|---|---|---|---|---|---|---|---|---|---|---|---|---|
| | $\Delta^{+}_{1\leftarrow 2}$ | $\Delta^{-}_{1\leftarrow 2}$ | $\Delta^{+}_{2\leftarrow 1}$ | $\Delta^{-}_{2\leftarrow 1}$ | $\Delta^{+}_{1\leftarrow 2}$ | $\Delta^{-}_{1\leftarrow 2}$ | $\Delta^{+}_{2\leftarrow 1}$ | $\Delta^{-}_{2\leftarrow 1}$ | $\Delta^{+}_{1\leftarrow 2}$ | $\Delta^{-}_{1\leftarrow 2}$ | $\Delta^{+}_{2\leftarrow 1}$ | $\Delta^{-}_{2\leftarrow 1}$ | $\Delta^{+}_{1\leftarrow 2}$ | $\Delta^{-}_{1\leftarrow 2}$ | $\Delta^{+}_{2\leftarrow 1}$ | $\Delta^{-}_{2\leftarrow 1}$ | $\Delta^{+}_{1\leftarrow 2}$ | $\Delta^{-}_{1\leftarrow 2}$ | $\Delta^{+}_{2\leftarrow 1}$ | $\Delta^{-}_{2\leftarrow 1}$ | $\Delta^{+}_{1\leftarrow 2}$ | $\Delta^{-}_{1\leftarrow 2}$ | $\Delta^{+}_{2\leftarrow 1}$ | $\Delta^{-}_{2\leftarrow 1}$ | $\Delta^{+}_{1\leftarrow 2}$ | $\Delta^{-}_{1\leftarrow 2}$ | $\Delta^{+}_{2\leftarrow 1}$ | $\Delta^{-}_{2\leftarrow 1}$ |
| **Av** | 0.721 | -0.446 | 0.801 | -0.946 | 0.571 | -0.449 | 0.601 | -0.706 | 0.531 | -0.463 | 0.562 | -0.640 | 0.539 | -0.400 | 0.514 | -0.677 | 0.579 | -0.459 | 0.739 | -0.849 | 0.084 | -0.065 | 0.011 | -0.090 | 0.248 | -0.120 | 0.052 | -0.228 |
| **Smu** | | | | | 0.145 | -0.029 | 0.022 | -0.706 | 0.086 | -0.030 | 0.021 | -0.640 | 0.040 | -0.027 | 0.020 | -0.677 | 0.021 | -0.946 | 0.207 | -0.019 | 0.818 | -0.946 | 0.233 | -0.078 | 0.796 | -0.946 | 0.441 | -0.192 |
| **Smi** | | | | | | | | | 0.015 | -0.706 | 0.100 | -0.014 | 0.018 | -0.706 | 0.241 | -0.018 | 0.020 | -0.706 | 0.163 | -0.024 | 0.558 | -0.706 | 0.300 | -0.071 | 0.640 | -0.631 | 0.204 | -0.228 |
| **Sp** | | | | | | | | | | | | | 0.232 | -0.017 | 0.018 | -0.677 | 0.018 | -0.640 | 0.064 | -0.024 | 0.529 | -0.640 | 0.245 | -0.074 | 0.652 | -0.640 | 0.267 | -0.228 |
| **Sl** | | | | | | | | | | | | | | | | | 0.021 | -0.677 | 0.227 | -0.027 | 0.562 | -0.677 | 0.275 | -0.075 | 0.587 | -0.677 | 0.381 | -0.197 |
| **Ss** | | | | | | | | | | | | | | | | | | | | | 0.743 | -0.849 | 0.257 | -0.079 | 0.755 | -0.737 | 0.198 | -0.228 |
| **Bf** | | | | | | | | | | | | | | | | | | | | | | | | | 0.253 | -0.035 | 0.090 | -0.228 |

**Table S5. Growth rates of species in pairwise cocultures calculated using MICOM.** Subscript 1 refers to the species in the row and 2 to the column. Units are $h^{-1}$.

| | Smu | | Smi | | Sp | | Sl | | Ss | | Bf | | Lc | |
|---|---|---|---|---|---|---|---|---|---|---|---|---|---|---|
| | $\mu_1^c$ | $\mu_2^c$ | $\mu_1^c$ | $\mu_2^c$ | $\mu_1^c$ | $\mu_2^c$ | $\mu_1^c$ | $\mu_2^c$ | $\mu_1^c$ | $\mu_2^c$ | $\mu_1^c$ | $\mu_2^c$ | $\mu_1^c$ | $\mu_2^c$ |
| **Av** | 0.887 | 0.822 | 0.741 | 0.552 | 0.696 | 0.549 | 0.923 | 0.959 | 0.832 | 0.725 | 0.466 | 0.657 | 0.436 | 0.356 |
| **Smu** | | | 0.552 | 0.543 | 0.522 | 0.509 | 0.501 | 0.495 | 0.535 | 0.539 | 0.869 | 0.519 | 0.742 | 0.653 |
| **Smi** | | | | | 0.070 | 0.661 | 0.454 | 0.453 | 0.475 | 0.515 | 0.572 | 0.600 | 0.559 | 0.585 |
| **Sp** | | | | | | | 0.441 | 0.438 | 0.422 | 0.483 | 0.604 | 0.333 | 0.698 | 0.714 |
| **Sl** | | | | | | | | | 0.520 | 0.530 | 0.890 | 0.753 | 0.318 | 0.755 |
| **Ss** | | | | | | | | | | | 0.769 | 0.414 | 0.847 | 0.800 |
| **Bf** | | | | | | | | | | | | | 0.275 | 0.268 |

**Table S6. Net interactions between species estimated based on growth rates calculated using MICOM.** Subscript 1 refers to the species in the row and 2 to the column. Units are $h^{-1}$.

| | Smu | | Smi | | Sp | | Sl | | Ss | | Bf | | Lc | |
|---|---|---|---|---|---|---|---|---|---|---|---|---|---|---|
| | $\Delta_{1\leftarrow 2}^{net}$ | $\Delta_{2\leftarrow 1}^{net}$ | $\Delta_{1\leftarrow 2}^{net}$ | $\Delta_{2\leftarrow 1}^{net}$ | $\Delta_{1\leftarrow 2}^{net}$ | $\Delta_{2\leftarrow 1}^{net}$ | $\Delta_{1\leftarrow 2}^{net}$ | $\Delta_{2\leftarrow 1}^{net}$ | $\Delta_{1\leftarrow 2}^{net}$ | $\Delta_{2\leftarrow 1}^{net}$ | $\Delta_{1\leftarrow 2}^{net}$ | $\Delta_{2\leftarrow 1}^{net}$ | $\Delta_{1\leftarrow 2}^{net}$ | $\Delta_{2\leftarrow 1}^{net}$ |
| **Av** | 0.365 | -0.114 | 0.220 | -0.146 | 0.174 | -0.085 | 0.401 | 0.289 | 0.310 | -0.116 | -0.055 | 0.568 | -0.086 | 0.130 |
| **Smu** | | | -0.384 | -0.155 | -0.415 | -0.125 | -0.436 | -0.175 | -0.402 | -0.302 | -0.068 | 0.430 | -0.195 | 0.428 |
| **Smi** | | | | | -0.628 | 0.027 | -0.245 | -0.217 | -0.223 | -0.325 | -0.127 | 0.511 | -0.139 | 0.359 |
| **Sp** | | | | | | | -0.193 | -0.232 | -0.211 | -0.358 | -0.029 | 0.244 | 0.064 | 0.489 |
| **Sl** | | | | | | | | | -0.150 | -0.311 | 0.220 | 0.664 | -0.353 | 0.530 |
| **Ss** | | | | | | | | | | | -0.071 | 0.325 | 0.006 | 0.574 |
| **Bf** | | | | | | | | | | | | | 0.186 | 0.042 |

**Table S7. The quartets of the interactions between species estimated based on growth rates calculated using MICOM.** Subscript 1 refers to the species in the row and 2 to the column. Units are $h^{-1}$.

| | Smu | | | | Smi | | | | Sp | | | | Sl | | | | Ss | | | | Bf | | | | Lc | | | |
|---|---|---|---|---|---|---|---|---|---|---|---|---|---|---|---|---|---|---|---|---|---|---|---|---|---|---|---|---|
| | $\Delta^{+}_{1\leftarrow2}$ | $\Delta^{-}_{1\leftarrow2}$ | $\Delta^{+}_{2\leftarrow1}$ | $\Delta^{-}_{2\leftarrow1}$ | $\Delta^{+}_{1\leftarrow2}$ | $\Delta^{-}_{1\leftarrow2}$ | $\Delta^{+}_{2\leftarrow1}$ | $\Delta^{-}_{2\leftarrow1}$ | $\Delta^{+}_{1\leftarrow2}$ | $\Delta^{-}_{1\leftarrow2}$ | $\Delta^{+}_{2\leftarrow1}$ | $\Delta^{-}_{2\leftarrow1}$ | $\Delta^{+}_{1\leftarrow2}$ | $\Delta^{-}_{1\leftarrow2}$ | $\Delta^{+}_{2\leftarrow1}$ | $\Delta^{-}_{2\leftarrow1}$ | $\Delta^{+}_{1\leftarrow2}$ | $\Delta^{-}_{1\leftarrow2}$ | $\Delta^{+}_{2\leftarrow1}$ | $\Delta^{-}_{2\leftarrow1}$ | $\Delta^{+}_{1\leftarrow2}$ | $\Delta^{-}_{1\leftarrow2}$ | $\Delta^{+}_{2\leftarrow1}$ | $\Delta^{-}_{2\leftarrow1}$ | $\Delta^{+}_{1\leftarrow2}$ | $\Delta^{-}_{1\leftarrow2}$ | $\Delta^{+}_{2\leftarrow1}$ | $\Delta^{-}_{2\leftarrow1}$ |
| **Av** | 0.824 | -0.458 | 0.822 | -0.937 | 0.630 | -0.411 | 0.551 | -0.698 | 0.633 | -0.458 | 0.549 | -0.634 | 0.923 | -0.522 | 0.959 | -0.670 | 0.753 | -0.443 | 0.725 | -0.841 | 0.466 | -0.522 | 0.649 | -0.081 | 0.436 | -0.522 | 0.321 | -0.191 |
| **Smu** | | | | | 0.354 | -0.738 | 0.247 | -0.402 | 0.323 | -0.738 | 0.221 | -0.346 | 0.245 | -0.681 | 0.190 | -0.366 | 0.195 | -0.596 | 0.182 | -0.484 | 0.868 | -0.937 | 0.513 | -0.084 | 0.742 | -0.937 | 0.605 | -0.178 |
| **Smi** | | | | | | | | | 0.069 | -0.698 | 0.085 | -0.057 | 0.221 | -0.466 | 0.209 | -0.426 | 0.200 | -0.423 | 0.235 | -0.561 | 0.572 | -0.698 | 0.583 | -0.073 | 0.559 | -0.698 | 0.537 | -0.178 |
| **Sp** | | | | | | | | | | | | | 0.230 | -0.422 | 0.235 | -0.467 | 0.154 | -0.365 | 0.203 | -0.561 | 0.604 | -0.634 | 0.327 | -0.083 | 0.698 | -0.634 | 0.714 | -0.226 |
| **Sl** | | | | | | | | | | | | | | | | | 0.276 | -0.426 | 0.351 | -0.663 | 0.890 | -0.670 | 0.753 | -0.089 | 0.318 | -0.670 | 0.755 | -0.226 |
| **Ss** | | | | | | | | | | | | | | | | | | | | | 0.769 | -0.841 | 0.406 | -0.081 | 0.847 | -0.841 | 0.800 | -0.226 |
| **Bf** | | | | | | | | | | | | | | | | | | | | | | | | | 0.275 | -0.088 | 0.268 | -0.226 |
